# Base-resolution mapping reveals distinct m^1^A methylome in nuclear- and mitochondrial-encoded transcripts

**DOI:** 10.1101/202747

**Authors:** Xiaoyu Li, Xushen Xiong, Meiling Zhang, Kun Wang, Ying Chen, Jun Zhou, Yuanhui Mao, Jia Lv, Danyang Yi, Xiao-Wei Chen, Chu Wang, Shu-Bing Qian, Chengqi Yi

**Affiliations:** State Key Laboratory of Protein and Plant Gene Research, School of Life Sciences, Peking University, Beijing 100871, China.; Academy for Advanced Interdisciplinary Studies, Peking University, Beijing 100871, PR China; Department of Chemical Biology and Synthetic and Functional Biomolecules Center, College of Chemistry and Molecular Engineering, Peking University, Beijing 100871, China.; Division of Nutritional Sciences, Cornell University, Ithaca, New York 14853, USA; Institute of Molecular Medicine, Peking University, Beijing, China; Peking-Tsinghua Center for Life Sciences, Peking University, Beijing, China.

## Abstract

Gene expression can be post-transcriptionally regulated via dynamic and reversible RNA modifications. *N*^1^-methyladenosine (m^1^A) is a recently identified mRNA modification; however, little is known about its precise location, regulation and function. Here, we develop a base-resolution m^1^A profiling method, based on m^1^A-induced misincorporation during reverse transcription, and report distinct classes of m^1^A methylome in the human transcriptome. m^1^A in 5’-UTR, particularly those at the first nucleotide of mRNA, associate with increased translation efficiency. A different subset of m^1^A exhibit a GUUCRA tRNA-like motif, are evenly distributed in the transcriptome and are dependent on the methyltransferase TRMT6/61A. Additionally, we show for the first time that m^1^A is prevalent in the mitochondrial-encoded transcripts. Manipulation of m^1^A level via TRMT61B, a mitochondria-localizing m^1^A methyltransferase, demonstrates that m^1^A in mitochondrial mRNA interferes with translation. Collectively, our approaches reveal distinct classes of m^1^A methylome and provide a resource for functional studies of m^1^A-mediated epitranscriptomic regulation.

## INTRODUCTION

More than 100 different types of post-transcriptional modifications have been identified so far (Machnicka et al., 2013). Recent breakthroughs in sequencing technologies have greatly advanced our understanding to the location, regulation, and function of RNA modifications in the transcriptome (Frye et al., 2016; Fu et al., 2014; Helm and Motorin, 2017; Li et al., 2016b), leading to the emerging field of epitranscriptomics (He, 2010; Saletore et al., 2012). One such example is N^1^-methyladenosine (m^1^A), a prevalent modification in non-coding RNA (ncRNA) and messenger RNA (mRNA) (Anderson and Droogmans, 2005; Roundtree et al., 2017). m^1^A was first documented more than 50 years ago (Dunn, 1961); later it was found to be a primordial RNA modification across the three major phylogenetic domains (Machnicka et al., 2013). In human cells, m^1^A is found at position 9 and 58 of human mitochondrial and cytoplasmic tRNAs, catalyzed by TRMT10C, TRMT61B and TRMT6/61A, respectively (Chujo and Suzuki, 2012; Ozanick et al., 2005; Vilardo et al., 2012); it is also present at position 1322 of 28S rRNA, catalyzed by NML (Waku et al., 2016). Its unique physicochemical property has also endowed m^1^A with pivotal roles in maintaining the proper structure and function of these ncRNAs (Roundtree et al., 2017). m^1^A in tRNA has also been systematically evaluated by microarray and sequencing (Cozen et al., 2015; Saikia et al., 2010; Zheng et al., 2015); more recently, m^1^A58 is shown to be reversible by ALKBH1, demonstrating an example of reversible tRNA modification in translation regulation (Liu et al., 2016). In addition to ncRNAs, m^1^A is also found to be a dynamic modification in mammalian mRNA, with strong enrichment in the 5’-UTR (Dominissini et al., 2016; Li et al., 2016a).

Despite such rapid progress, a high-resolution profile of the mammalian m^1^A methylome is still lacking, significantly limiting our understanding and functional characterization of this newly discovered mRNA modification. Previous m^1^A profiling technologies have a resolution of about tens of nucleotides to several hundred nucleotides, primarily determined by the size of RNA fragments in these experiments (Dominissini et al., 2016; Li et al., 2016a). In addition, the methyltransferase(s) and functional consequence of mRNA m^1^A modification is poorly understood. Hence, except for a handful positions in rRNA and tRNA, little is known about the precise location, regulation and function of m^1^A in the human transcriptome.

Here, we report a base-resolution method to profile m^1^A in the human transcriptome. Our method is based on m^1^A-induced misincorporation during reverse transcription and reveals distinct classes of m^1^A methylome: a major group of m^1^A sites that are enriched in 5’-UTR, a small subset of GUUCRA(“R” denotes a purine)-tRNA like m^1^A sites with relatively even distribution in the transcriptome, and prevalent m^1^A modification in the CDS of 10/13 mitochondrial(mt)-encoded transcripts. m^1^A sites in the 5’-UTR, particularly those located at the first nucleotide of mRNA transcripts (or “cap+1” position), are associated with increased translation efficiency. In contrast, m^1^A in the CDS of mt-mRNA inhibits translation. Collectively, our approaches reveal distinct classes of base-resolution m^1^A methylome in the nuclear- and mitochondrial-encoded transcripts, and provide an in-depth resource towards elucidating the functions of m^1^A methylation in mRNA.

## RESULTS

### m^1^A-induced misincorporation during reverse transcription

Because m^1^A can cause both truncation and misincorporation during cDNA synthesis (Hauenschild et al., 2015; Zubradt et al., 2017), we first established the truncation and mutation profiles of different reserve transcriptases (RTases). We systematically compared the performance of several commercially available RTases (including AMV, SuperScript II, SuperScript III and TGIRT) under different conditions (Figure S1). We found that m^1^A can precisely induce misincorporation at the site of modification, while m^1^A-induced truncation is less accurate and can occur to the neighboring nucleotides. In addition, the truncation profile could be complicated by RNA secondary structures and the fragmentation process needed for library preparation. We concluded that the mutation profile contains a higher signal/noise ratio and is more precise in detecting the exact position of m^1^A. Among the RTases we tested, TGIRT demonstrated excellent read-through efficiency and relatively high mutation frequency at the site of m^1^A (Figure S1B), consistent with the recent DMS-MaPseq and DM-tRNA-seq results (Zheng et al., 2015; Zubradt et al., 2017). Moreover, we employed a ligation-based strand-specific library preparation protocol (Van Nostrand et al., 2016), which ensures that the m^1^A-induced mutation is within the sequenced fragment (see Method Details).

Because we only observed ~40-50% mutation rate at m^1^A1322 in 28S rRNA (Figure S2A), which is known to be of high modification level, we further examined the quantitative capability of TGIRT. We chemically synthesized two model RNA sequences with site-specific m^1^A modification. For m^1^A sites with ~97-98% modification level (measured by quantitative mass spectrometry) (Figure S2B), we consistently observed ~66-75% misincorporation (Figure S2C); the mismatch rate dropped non-linearly when we gradually lowered the modification level. Even with ~50% m^1^A modification, a mismatch rate of only ~9-10% was observed (Figure S2C). These findings suggest that the observed mutation rate is an underestimation of the actual modification level. While the TGIRT-based procedure can still detect m^1^A sites of high modification level, sequencing RNA directly with TGIRT may not be able to capture the m^1^A sites with averaging modification level in the transcriptome (~20% as previously measured by microarray) (Dominissini et al., 2016). To improve the sensitivity for transcriptome-wide m^1^A detection, we decided to couple the TGIRT-based procedure with a pre-enrichment step and an additional *in vitro* demethylation step (Figure 1A). We first show that *in vitro* demethylation reaction mediated by the demethylase AlkB is more efficient than the Dimroth reaction, demonstrating ~98% and ~80% efficiency (Figure S2D), respectively. In addition, the extended treatment of RNA in alkaline condition during the Dimroth reaction leads to excessive RNA degradation (Figure S2E), potentially causing loss of RNA molecules. By integrating the enrichment and demethylation steps, we successfully maximized the dynamic range of m^1^A-induced mutational signature for m^1^A1322 in 28S rRNA (~47%, ~95% and ~0.9% in the input, (−) and (+) demethylase samples, respectively), allowing sensitive and confident m^1^A detection (Figure 1B). We termed our approach misincorporation-assisted profiling of m^1^A, or m^1^A-MAP.

**Figure 1.**
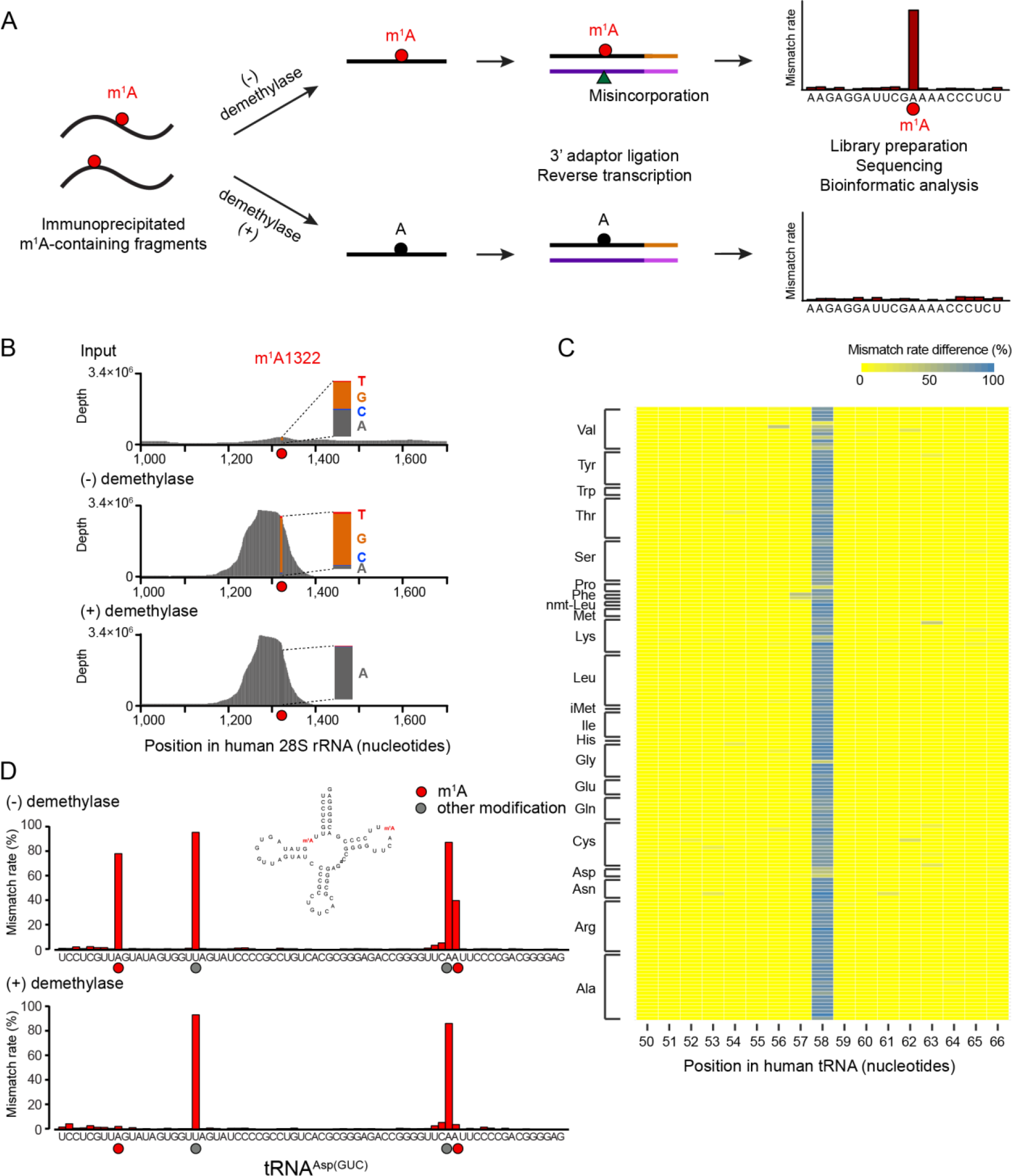
m^1^A-MAP utilizes m^1^A-induced misincorporation to detect m^1^A sites at singlenucleotide resolution. (**A**) Scheme of m^1^A-MAP. We optimized the conditions of RT so as to allow efficient misincorporation in cDNA synthesis. The use of an m^1^A antibody pre-enriches the m^1^A-containing RNA fragments, thereby maximizing the misincorporation signal; and the use of demethylase treatment improves the confidence of detection. An m^1^A modification is called depending on the difference and fold change of mismatch rate between the (−) and (+) demethylase samples (see Method Details). (**B**) m^1^A-MAP maximizes the misincorporation signal for m^1^A1322 on 28S rRNA. (**C**) m^1^A-MAP detects m^1^A58 for the cytosolic tRNAs. Shown here is the difference of mismatch rate between the (−) and (+) demethylase samples. (**D**) m^1^A-MAP detects a novel m^1^A site at position 9 in the cytosolic human tRNA^Asp(GUC)^. The mismatch rate for m^1^A58 is also reduced after demethylase treatment, while two other modifications (at position 20 and 57) are not sensitive to demethylation, representing other types of RNA modifications. See also Figure S1 and S2.

### m^1^A-MAP detects known and novel m^1^A in tRNA

We next applied m^1^A-MAP to tRNA. In mammals, m^1^A can occur at position 9, 14 and 58 of tRNA(Anderson and Droogmans, 2005). m^1^A14 has been reported only in tRNA^Phe^ and is considered to be very rare (Machnicka et al., 2013); we did not observe any m^1^A modification at position 14 for cytosolic tRNAs in HEK293T cells (Table S1). m^1^A58 is conserved across the three domains of life; previous tRNA microarray and sequencing data has reported hypomodified tRNAs at this position (Cozen et al., 2015; Saikia et al., 2010; Zheng et al., 2015). Our results confirmed that m^1^A58 is globally present in the cytosolic tRNAs (Figure 1C and S2F). The m^1^A9 modification exists only in archaea tRNA or mammalian mitochondrial tRNA; interestingly, we observed a novel m^1^A9 site for cytosolic tRNA^Asp(GUC)^, representing the first cytosolic tRNA with m^1^A modification at position 9 (Figure 1D). Collectively, these observations suggest that m^1^A-MAP is highly sensitive in detecting m^1^A at single-base resolution.

### Single-nucleotide resolution m^1^A methylome in the transcriptome

We then sought to detect transcriptome-wide m^1^A methylome at single-base resolution. We defined two parameters to evaluate the m^1^A-MAP data: difference of mismatch rate and fold change of mismatch rate (see Method Details). To minimize the effect of mismatch rate variation during m^1^A identification, we rigorously tested our threshold and identified 740 m^1^A sites in the 293T transcriptome (Table S1-3). To evaluate potential false positives caused by m^1^A-independent mismatch, we performed the opposite calculation and retrieved only 17 such sites (see Method Details). Moreover, we also systematically evaluated the mutation pattern of the identified m^1^A sites in the transcriptome. Using m^1^A sites in tRNA as positive controls, we found that m^1^A-induced mutation is more strongly influenced by its 5’-nucleotide than the 3’-nucleotide; importantly, a similar sequence-dependent feature is also observed for m^1^A sites in mRNA (Figure 2A and S3A). Therefore, we conclude that our strict threshold allowed us to confidently detect transcriptome-wide m^1^A sites at single-nucleotide resolution.

**Figure 2.**
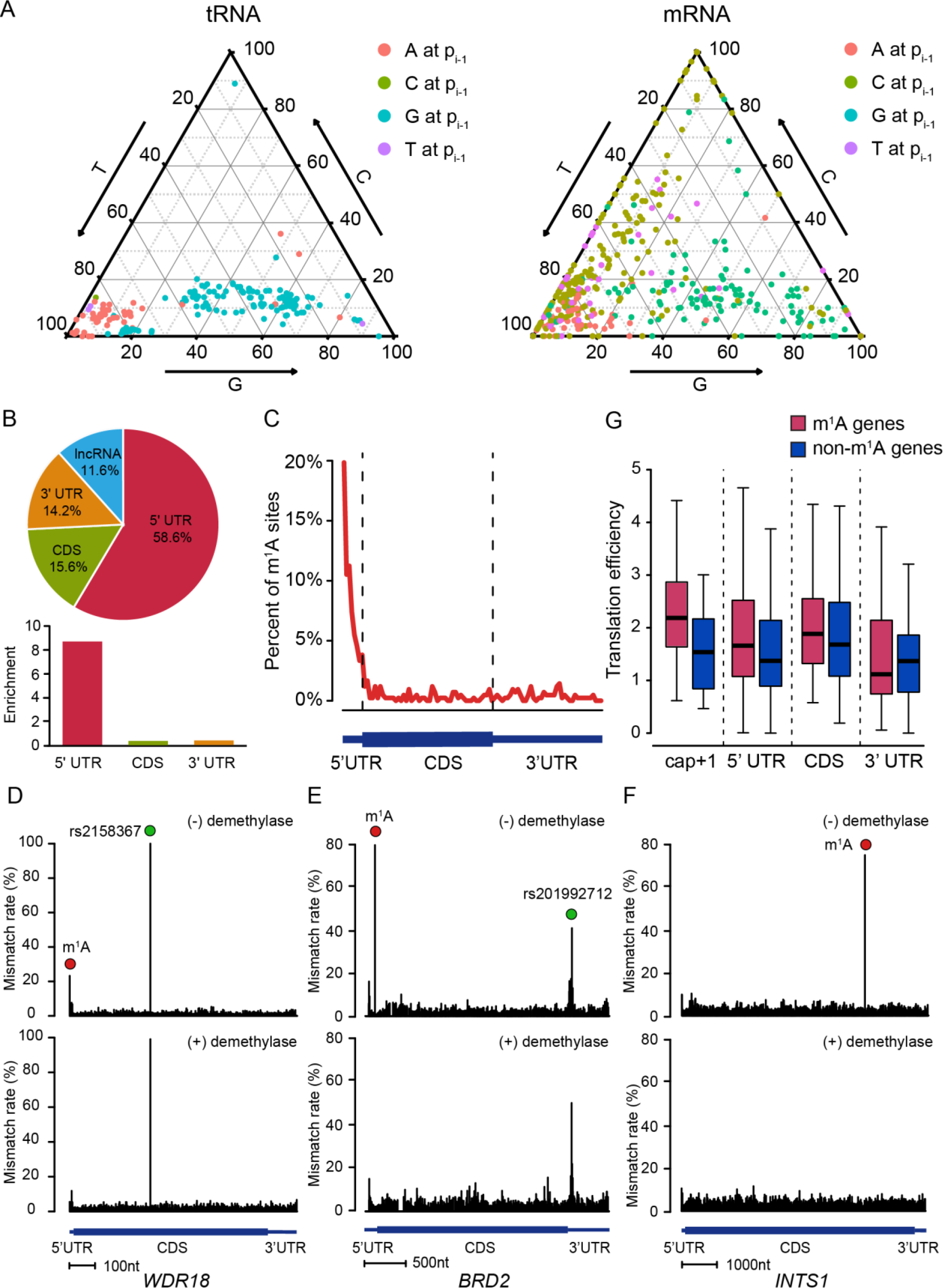
Single-nucleotide resolution m^1^A methylome in the human transcriptome. (**A**) Mutation pattern of m^1^A sites in mRNA resembles that in tRNA. Shown here is the sequence-dependent mutation profile of m^1^A sites with regard to the immediate 5’ nucleotide. (**B**) The pie chart shows the percentage of m^1^A sites in each non-overlapping segment. (**C**) Distribution of m^1^A sites across mRNA segments. Each segment was normalized according to its average length in Refseq annotation. (**D-F**) Representative views of a typical m^1^A site at the cap+1 position (the first transcribed nucleotide of mRNA) of *WDR*18 (**D**), the 5’-UTR of *BRD*2 (**E**), and the CDS of *INTS*1 mRNA (**F**). The demethylation-insensitive signals in (**D**) and (**E**), which are indicated as green dots, are known SNPs. The scale bars are indicated at the bottom of each panel. (**G**) m^1^A in the cap+1 position and 5’-UTR positively correlates with increased translation efficiency. Transcripts with comparable expression level but without m^1^A sites were chosen as the negative control for each category. See also Figure S3.

Out of the 740 m^1^A modifications in the transcriptome (Figure S3B), 473 sites are located in mRNA and lncRNA molecules (Figure S3C and Table S2). Majority of these sites are within the 5’-UTR (Figure 2B and 2C), consistent with the previous finding (Dominissini et al., 2016; Li et al., 2016a). Our single-base profile also reveals new features of the m^1^A methylome: for instance, we found 24 m^1^A methylation sites that are present exactly at the first nucleotide of the 5’ end of the transcripts (Figure 2D and Table S2). Because the first two nucleotides of the 5’ end of mRNA are known to contain ribose methylation, it is likely that these transcripts have an m^1^A_m_ modification at the cap+1 site. We also found 3 additional transcripts with m^1^A methylation at the second nucleotide, potentially giving rise to m^1^A_m_ at cap+2 as well. For m^1^A located in CDS, while we did not observe a tendency for codon position, we did notice a mild preference for codon types, with Arg(CGA) being the most frequently modified by m^1^A (Figure S3D). No m^1^A is detected for AUG start codons. Representative examples of m^1^A sites identified from different mRNA regions are shown (Figure 2D-F and S3E). Two additional sites of high mutation, which are insensitive to the demethylase treatment, also appeared in the *WDR*18 and *BRD*2 examples (Figure 2D and 2E). By referring to the SNP database, we found that these two positions belong to annotated SNP sites, demonstrating the robustness of our approach in distinguishing true m^1^A sites from false signals (SNP, other modifications and etc.).

Because m^1^A is enriched in the 5’-UTR, we examined whether m^1^A could be involved in translation regulation. We performed ribosome profiling and compared the translation efficiency for transcripts with or without m^1^A. We found that m^1^A within the 5’-UTR positively correlates with the translation efficiency of mRNA (Figure 2G). This positive correlation is even stronger for m^1^A at the cap+1 position, but is not observed for m^1^A located in CDS nor 3’-UTR. This observation hints that m^1^A within different regions of mRNA may have different biological functions.

### A subset of m^1^A sites demonstrate a GUUCRA consensus motif

An unbiased motif detection using DREME revealed that a subset of m^1^A (53 sites) are found within a strong GUUCRA sequence (Figure 3A). Interestingly, these sites demonstrate a very different distribution pattern: instead of being enriched in the 5’-UTR, these sites are evenly distributed in the transcriptome (Figure 3B). Because this motif is reminiscent of the m^1^A-containing TψC loop in tRNA, we hypothesized that the tRNA methyltransferase complex TRMT6/61A could be responsible for these mRNA m^1^A sites. We first performed direct m^1^A sequencing (without antibody enrichment) to RNA population below 200nt. We found that the m^1^A58 sites within the GUUCNA motif experienced a global decrease of mutation rate in the TRMT6/61A knock-down sample, which was not observed for m^1^A58 sites that do not confine to the motif (Figure 3C, S4A and S4B). This result suggests that TRMT6/61A-mediated m^1^A methylation is highly sequence-specific, consistent with evidence from crystal structures (Finer-Moore et al., 2015). We then analyzed the secondary structure for the 53 mRNA m^1^A sites, and found highly conserved structural features compared to the T-loop of tRNA (Figure S4C and S4D). We also picked 3 m^1^A sites (in CDS, 3’-UTR and lncRNA, respectively) and examined their response after TRMT6/61A knock-down (Figure 3D). Our locus-specific approach (see Method Details), which enabled us to interrogate these sites with high sequencing depth, unambiguously demonstrated a decrease in mismatch rate after TRMT6/61A knock-down (Figure 3D). As a comparison, a non-motif m^1^A site located in a different structural context demonstrated an unaltered modification status (Figure 3E). Taken together, these observations suggest that in addition to tRNA, TRMT6/61A is also responsible for a subset of m^1^A sites in mRNA.

**Figure 3.**
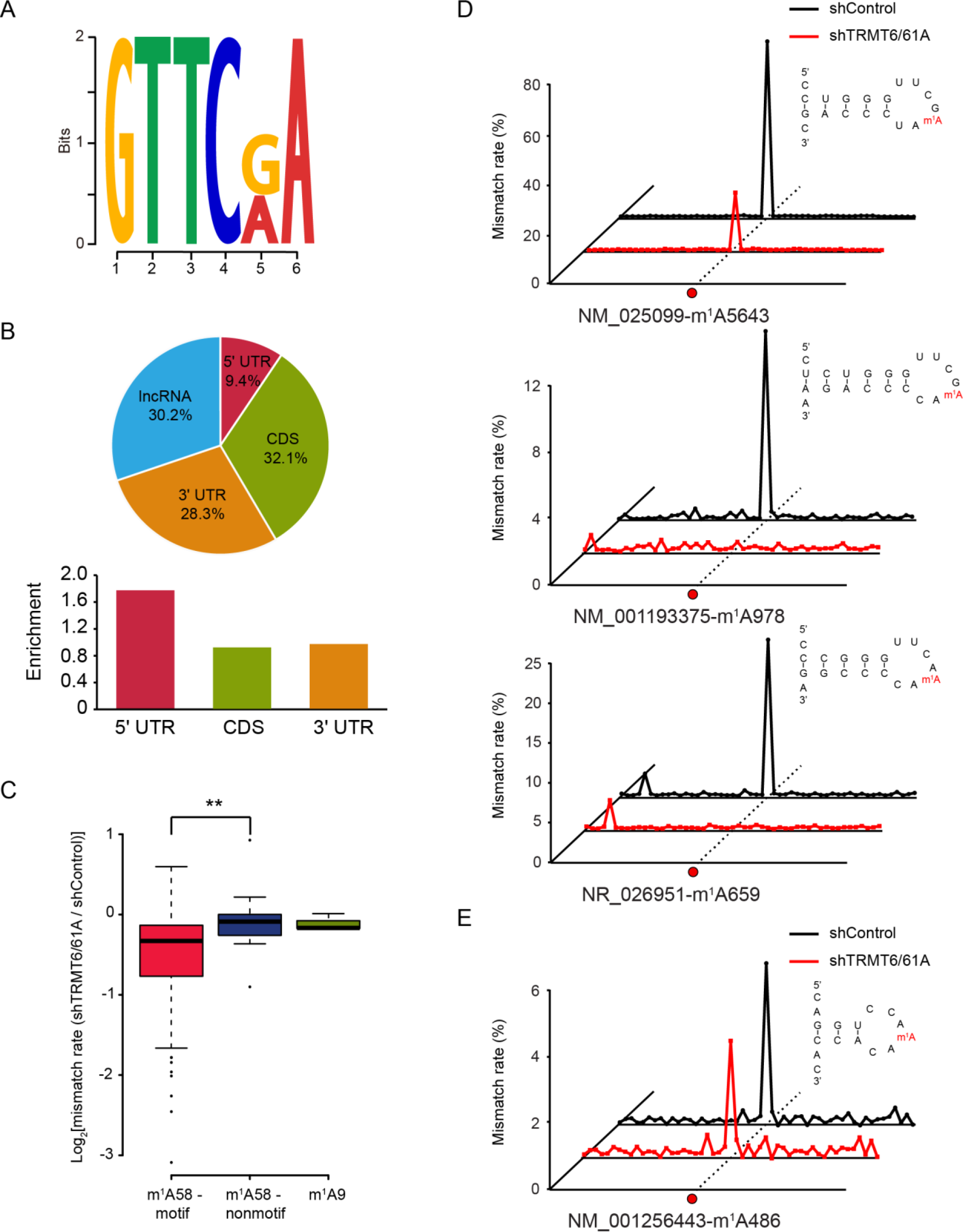
The tRNA methyltransferase complex TRMT6/61A catalyzes a subset of m1A methylation in mRNA. (**A**) methylation in mRNA.Motif analysis revealed a GUUCRA tRNA-like consensus for a group of m^1^A in mRNA, E-value=4.8e-009. (**B**) Pie chart showing the percentage of m^1^A sites in each non-overlapping segment. Comparing to the non-motif m^1^A sites in mRNA, these sites are evenly distributed in the transcriptome. (**C**) TRMT6/61A specifically targets m^1^A within the consensus sequence, while doesn't work on m^1^A in non-motif sequence of the T-loop nor m^1^A at the 9th position of tRNA, p-value <0.005. (**D**) Representative views of three mRNA targets of TRMT6/61A. The predicted RNA secondary structures are also shown, revealing a conserved stem-loop structure that harbors the GUUCRA motif. (**E**) An example of non-motif mRNA m^1^A site, showing similar mismatch rates in the control and TRMT6/61A knock-down samples. See also Figure S4.

### Distinct m^1^A methylome in the mitochondrial transcriptome

In addition to the nuclear-encoded transcripts, we also detected prevalent m^1^A modification in the mitochondrial (mt) transcriptome. mt-tRNAs are known to contain m^1^A at position 9 and 58 (Suzuki et al., 2011), catalyzed by TRMT10C and TRMT61B (Chujo and Suzuki, 2012; Vilardo et al., 2012), respectively. m^1^A-MAP showed that all the 14 mt-tRNAs bearing an adenosine residue at position 9 are m^1^A modified; for position 58, m^1^A was detected for the 3 known and 2 novel mt-tRNA molecules (Figure 4 and Table S3). For mt-rRNA, the only known m^1^A site is at position 947 of 16S rRNA (Bar-Yaacov et al., 2016). Interestingly, we additionally detected 7 and 10 novel m^1^A sites on 16S and 12S mt-rRNA, respectively (Figure 4 and Table S3). This is very different from cytosolic rRNA, where there is only one m^1^A site in 28S rRNA (m^1^A1322). Considering the length of these rRNA species, mt-rRNAs are much more heavily modified by m^1^A.

**Figure 4.**
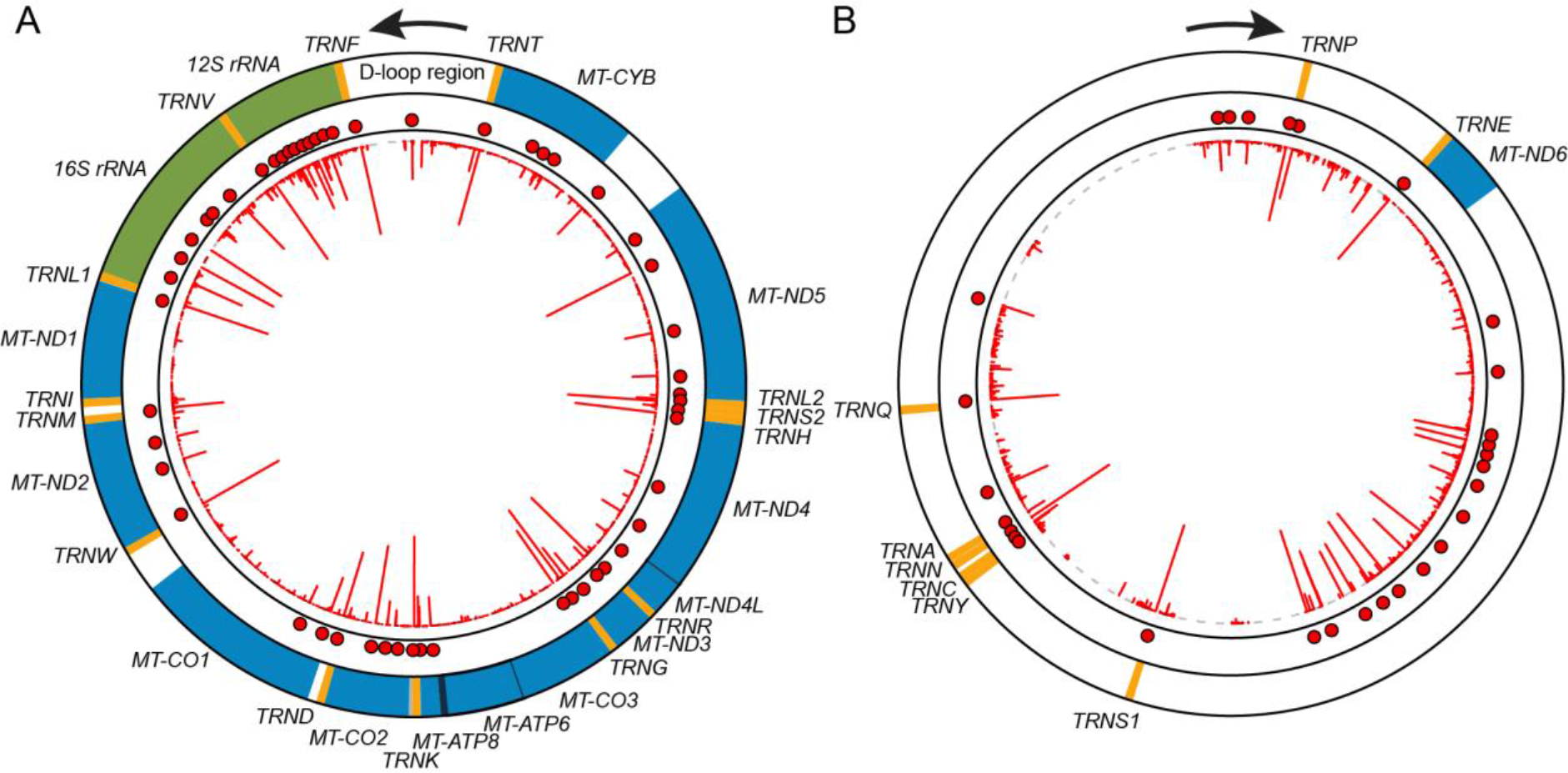
Distinct m^1^A methylome in the mitochondrial transcriptome. (**A**) m^1^A methylome of the heavy strand. Orange, green and blue colors represent tRNA, rRNA and mRNA, respectively. The red line in the inner circle represents the difference of mismatch rate of individual nucleotide while each red dot represents an identified m^1^A site. (**B**) m^1^A methylome of the light strand. See also Figure S5.

In human mitochondria, mRNAs are transcribed from the heavy and light strands as polycistronic units (Falkenberg et al., 2007; Mercer et al., 2011). The processed mt-mRNAs lack a cap at the 5’ end and contain no or short untranslated regions (Richter-Dennerlein et al., 2015; Rorbach and Minczuk, 2012; Temperley et al.,2010). We identified 22 m^1^A sites from 10/13 mitochondrial genes, in which 21 are residing in CDS and 1 is located in the 3’-UTR (Figure 4). This is distinct to the m^1^A methylome in the nuclear-encoded transcripts, where m^1^A is enriched in the 5’-UTR. In addition, no preference for codon types was observed, yet m^1^A appears to be more likely present at the third position of a codon in the CDS of mt-mRNA (Figure S5A). Moreover, we also identified 25 m^1^A sites within the intergenic spacers. 24/25 m^1^A sites are in the light strand (Figure 4B); some of these m^1^A sites could be within the 3’-UTR of *MT-ND6*, for which there is no current consensus of its 3’ end (Slomovic et al., 2005).

### m^1^A in mt-mRNA interferes with mitochondrial translation

We next sought to examine the biological consequence of m^1^A in the mt-mRNAs. Translation requires accurate base pairing between mRNA codons and the cognate tRNAs; however, m^1^A is known to block the canonical A:U base pairing. These facts prompted us to hypothesize that m^1^A in mt-mRNA, which are primarily located in CDS, could interfere with translation in mitochondria. We first integrated published mitochondria ribosome profiling data with m^1^A-MAP identified m^1^A methylome in mitochondria (Rooijers et al., 2013). We found a strong signal of mitochondrial ribosome stalling at the m^1^A site on *MT-ND5* (Figure 5A and S5B), whose modification level is the highest among all m^1^A sites in mt-mRNA. Due to the difficulty of an *in vitro* mitochondrial translation system (Smits et al., 2010), we sought to enzymatically manipulate the modification level of the mt-mRNAs. Two enzymes are known to introduce m^1^A in human mitochondria: TRMT10C generates m^1^A as well as m^1^G at position 9 in mitochondrial tRNAs, while TRMT61B is responsible for m^1^A at position 58 in some mitochondrial tRNAs and position 947 in 16S mt-rRNA (Bar-Yaacov et al., 2016; Chujo and Suzuki, 2012; Vilardo et al., 2012). Because TRMT10C is a subunit of the mitochondrial RNase P complex and is not specific for adenosine, we focused on TRMT61B. We utilized a qPCR-based assay to quantitatively evaluate the modification status of the m^1^A sites in mt-mRNAs (see Method Details); we found that while TRMT61B knock-down mildly reduced the m^1^A level (Figure S5C and S5D), TRMT61B overexpression greatly increased the m^1^A modification level of several mt-mRNAs (Figure 5B and S5D). This observation suggests that in addition to mt-tRNA and mt-rRNA, TRMT61B could also target mt-mRNA. Because of the high efficiency of TRMT61B overexpression in increasing the m^1^A level, we used mass spectrometry to quantitatively measure the mitochondrial protein level upon TRMT61B overexpression (Figure S5E). Indeed, TRMT61B overexpression led to a reduced protein level for *MT-CO*2 and *MT-CO*3 (Figure 5C and S5F), which are targets of TRMT61B. We further confirmed this observation for the MT-CO2 protein using Western blot (Figure 5D). Collectively, these results suggest that m^1^A in mt-mRNA interferes with translation in mitochondria.

**Figure 5.**
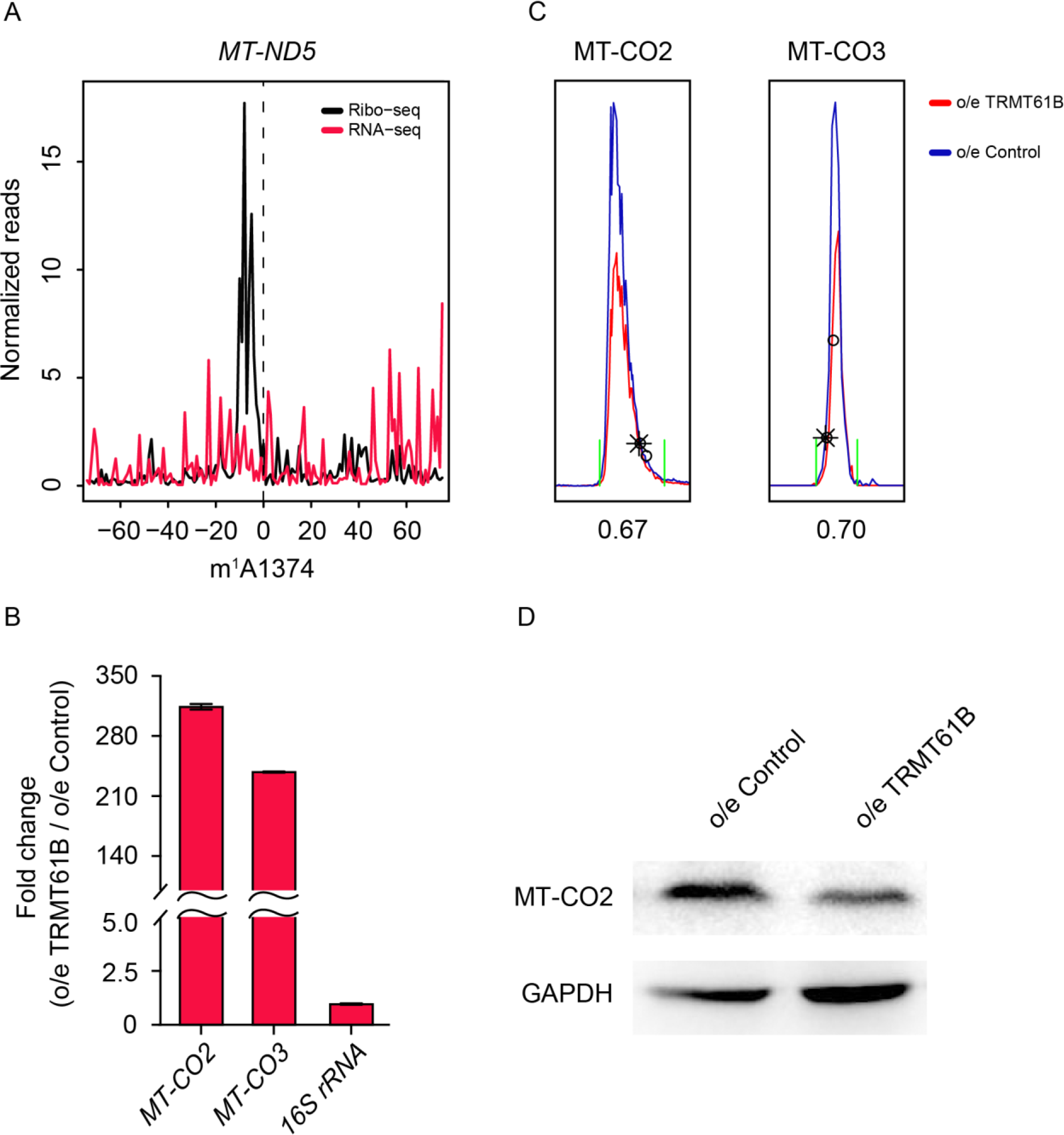
m^1^A in the mitochondrial mRNA interferes with translation. (**A**) Mitochondrial ribosome stalling at the m^1^A site of the *MT-ND5* mRNA. The density of the 5’ end of footprints was calculated for each position surrounding the m^1^A site. Ribosome profiling and RNA-seq data was taken from a published study (see Method Details). (**B**) TRMT61B overexpression led to increased m^1^A level in *MT-CO2* and *MT-CO3* mRNA, as measured by the qPCR-based assay. Data are mean ± SD; n = 2. (**C**) Extracted ion chromatograms of MT-CO2 and MT-CO3, showing decreased protein level. (**D**) Western blot of MT-CO2 upon TRMT61B overexpression. See also Figure S5.

## DISCUSSION

In this study, we developed a single-nucleotide resolution method for transcriptome-wide identification of m^1^A in human cells. m^1^A-MAP utilizes the m^1^A-induced misincorporation in cDNA synthesis to achieve base-resolution m^1^A detection. This enabled us to identify m^1^A modification not only at the mRNA cap, but also within a GUUCRA tRNA-like sequence motif. In principle, such misincorporation-dependent strategy could be applied to estimate the modification status of RNA sites of interest. However, our results in both rRNA and model RNA sequences strongly suggest that such estimation should be done with caution: TGIRT underestimates m^1^A modification level and m^1^A-induced mismatch decreases in a non-linear fashion as the modification level decreases. Additionally, the sequence context of RNA has also been reported to affect the mutation rate (Hauenschild et al., 2015; Zubradt et al., 2017). Hence, while direct sequencing (without enrichment) could still detect m^1^A sites with high modification level, m^1^A sites with averaging modification level or those located within a non-optimal context for mismatch induction could be missed. Therefore, coupling the pre-enrichment step to the mutational signature is necessary to improve the detection sensitivity. In addition, to achieve high confidence, we employed an *in vitro* demethylation step, which enabled us to distinguish true m^1^A sites from false signals (SNP, other modifications, and etc.) in the transcriptome. The combined use of mutational signature, the pre-enrichment step and the *in vitro* demethylation step enabled m^1^A-MAP to achieve high sensitivity and confidence.

Our study revealed that two known m^1^A modification machinery, TRMT6/61A and TRMT61B, could work on mRNA as well. The hetero complex TRMT6/61A recognizes the sequence and structure of the tRNA T-loop and installs an m^1^A at the 58 position (Finer-Moore et al., 2015). Consistent with this knowledge, we found that the TRMT6/61A-dependent mRNA m^1^A sites are also confined to a hairpin structure mostly frequently with a 7nt loop, reminiscent of the tRNA T-loop. In contrast, we did not find an obvious sequence context for the m^1^A in mt-mRNA. In fact, TRMT61B appears to be a promiscuous enzyme that also modifies mt-tRNA and mt-rRNA; the substrate specificity and the underlying mechanism of TRMT61B in the human mitochondria remains to be determined. In addition, the fact that both TRMT6/61A and TRMT61B are known tRNA modification enzymes also reminds us of the modification machinery for other mRNA modifications. For instance, eukaryotic ψ synthases PUS1, PUS7 and TRUB1 can work on both tRNA and mRNA (Carlile et al., 2014; Li et al., 2015; Lovejoy et al., 2014; Safra et al., 2017; Schwartz et al., 2014a); yet the modification complex for m^6^A, consisting at least METTL3, METTL14, WTAP and KIAA1429, appears to be dedicated to mRNA (Bokar et al., 1997; Liu et al., 2014; Ping et al., 2014; Schwartz et al., 2014b). In the case of m^1^A, the enzyme(s) responsible for the majority of the modification sites in mRNA remains to be identified. It would be interesting to see if such machinery is specific for mRNA or promiscuous for multiple RNA substrates.

Our base-resolution m^1^A profiles reveal distinct m^1^A methylome in the human transcriptome. m^1^A is enriched in the 5’-UTR; and only 5’-UTR m^1^A sites, but not those in CDS or 3’-UTR, are correlated with higher translation efficiency. Different from the m^1^A sites in the nuclear-encoded transcripts, m^1^A in mt-mRNA are primarily located in CDS and inhibit translation. In addition, we also identified a notable group of m^1^A methylation adjacent to the mRNA cap, raising the possibility of m^1^A_m_ modification at the first position of RNA transcripts. A related but different modification, m^6^A_m_, is also known to be present at this position and is recently reported to improve mRNA stability (Mauer et al., 2017). While m^1^A-MAP could not discriminate m^1^A from m^1^A_m_, neither m^6^A nor m^6^A_m_ should induce misincorporation during reverse transcription, nor become sensitive to demethylation by AlkB. Hence, methylation to the N1 and N6 position of adenosine within the cap should represent distinct types of modifications.

Our results also revealed m^1^A in mt-mRNA for the first time and showed that such methylation interferes with translation. By manipulating the m^1^A level via TRMT61B, we monitored the corresponding changes of the mitochondrial protein level by both quantitative mass spectrometry and Western blot. TRMT61B modifies both mt-tRNA and mt-rRNA (Bar-Yaacov et al., 2016; Chujo and Suzuki, 2012); and these m^1^A sites are thought to be beneficial for their functions in translation. Because m^1^A sites in mt-mRNA have an opposing effect in translation, changes of protein synthesis level after TRMT61B knock-down could be a mixed result of contrary m^1^A sites in these different components (rRNA, tRNA and mRNA) of translation. Conversely, because16S rRNA and mt-tRNA^Leu(UUR)^ are already of high m^1^A level, we envisioned that TRMT61B overexpression should lead to a greater increase of m^1^A level in mRNA than in these ncRNAs. In this simplified scenario, we indeed observed greatly increased modification level for the mt-mRNA and detected reduced protein level for two mitochondrial proteins, whose mRNA transcripts are m^1^A modified. While our MS experiments detected the overall protein level, future experiments measuring the nascent protein level upon TRMT61B overexpression or knock-down could provide more detailed information regarding protein synthesis. In terms of the mechanism of m^1^A-mediated translational suppression, multiple possibilities need to be considered. For instance, due to the presence of a base-pairing interfering modification in the CDS, m^1^A could serve as a road block for the mitochondrial ribosome. In fact, recent *in vitro* translation experiments on synthetic RNA sequences have shown that m^1^A in CDS represses translation; the effect is stronger when m^1^A is at codon position 1 and 2, while m^1^A at the third position also mitigates translation (You et al., 2017). Thus, not only the density but also the exact position of m^1^A in mt-mRNA could influence protein synthesis. In addition, microRNA has been shown to enhance translation in mitochondria (Zhang et al., 2014). We analyzed the published CLASH results in which microRNA and their direct mRNA targets are captured (Helwak et al., 2013); interestingly, we found two m^1^A sites that are located within the experimentally verified targets of microRNA (Figure S5G). In fact, these m^1^A sites reside exactly within mRNA sequences that form base-pairing with the seed regions of microRNAs. More m^1^A sites in mt-mRNA were found within the predicted mt-mRNA targets of the microRNA seed regions (Figure S5H). While both the speculated mechanisms point to a suppressive role of m^1^A in mitochondria translation, alternative hypothesis and mechanism should also be tested in future experiments. Nevertheless, our discovery that m^1^A in mt-mRNA interferes with translation improves our understanding of translation regulation in human mitochondria.

In summary, our study demonstrated distinct classes of m^1^A methylome in the nuclear- and mitochondrial-encoded transcripts. Our single-nucleotide resolution m^1^A technology allowed the comprehensive profiling of m^1^A in the human transcriptome, providing a reference and resource for future investigations to elucidate the biological functions and mechanisms of this new epitranscriptomic mark.

## AUTHOR CONTRIBUTIONS

X.L., X.X., and C.Y. conceived the base-resolution method and performed sequencing; Y.C., M.Z., K.W., J.L., and D.Y. performed MS analysis and cell biology experiments, under the guidance of C.Y., C.W., and X.-W.C. X.L., K.W., and J.Z. performed ribosome profiling and translation experiments. X.X. designed and performed the bioinformatics analysis, with the help of Y.M. C.Y. and S.-B.Q. supervised the project. All authors commented on and approved the paper.

## ACKNOWLEDGMENTS

The authors would like to thank Prof. Ying Liu for advice on methyltransferase, Prof. Yongliang Zhao for sharing antibody, and Drs. Hui Li and Guilan Li for measurements with LC-MS/MS. This work was supported by the National Basic Research Foundation of China (nos. 2016YFC0900301 and 2014CB964900 to C.Y.), the National Natural Science Foundation of China (nos. 21522201 and 21472009 to C.Y.), US National Institutes of Health (R01 AG042400 and R01 GM122814 to S.-B.Q) and HHMI Faculty Scholar (55108556 to S.-B.Q).

## METHOD DETAILS

### Cell Culture and Antibodies

HEK293T (ATCC,CRL-11268) was used in this study and maintained in DMEM medium (Gibco) supplemented with 10% FBS (Gibco) and 1% penicillin/streptomycin (Gibco). Monoclonal mouse anti-m1A antibody was purchased from MBL (D345-3). Polyclonal rabbit anti-TRMT6 antibody was purchased from Santa Cruz Biotechnology (sc-271752). MT-CO2 antibody was purchased from Proteintech Group (55070-1-AP). Monoclonal mouse anti-β-Actin antibody was purchased from CWBiotech (CW0096). Monoclonal mouse anti-GAPDH antibody was purchased from CWBiotech (CW0100M).The secondary antibodies used are anti-mouse-IgG-HRP (CW0102; CWBiotech) and anti-rabbit-IgG-HRP (CW0103; CWBiotech).

### RNA isolation

Total RNA was isolated from cells using TRIzol, according to the manufacturer’s instructions (Invitrogen). An additional DNaseI treatment step was performed to avoid DNA contamination. For polyA^+^ RNA isolation, small RNA was depleted first using MEGAclear™ Transcription CleanUp Kit (Ambion), followed by two successive rounds of polyA^+^ selection using oligo(dT)_25_ dynabeads (Invitrogen). For small RNA isolation, RNA smaller than 200 nt were recovered from the flow-through fraction in the small RNA depletion step by ethanol precipitation.

### shRNA knock down of TRMT6/61A

The oligoes targeting TRMT6 and TRMT61A were annealed and cloned into the pLKO vector according to the TRC shRNA library protocol (http://www.broadinstitute.org/rnai/public/), respectively. The oligo sequences were listed below: TRMT6-FWD: CCGGGGGAAAGTT CTGAGT ATTTATCT CGAGATAAAT ACT CAGAACTTTCCCTTTTT G; TRMT6-RVS: AATTCAAAAAGGGAAAGTTCTGAGTATTTATCTCGAGATAAATACTCAGAACTTTCCC; TRMT61A-FWD: CCGGGAGGCCAGAGGCACCTTATATCTCGAGATATAAGGTGCCTCTGGCCTCTTTTTG; TRMT61A-RVS: AATTCAAAAAGAGGCCAGAGGCACCTTATATCTCGAGATATAAGGTGCCTCTGGCCTC. A scrambled shRNA was used as the mock control. Lentiviruses were packaged by co-transfecting HEK293T cells with pLKO-TRMT6, pLKO-TRMT61A, pCMV-dR8.91 and VSV-G plasmids, following the instructions from Broad Institute. The supernatants from transfected cells were harvested after 2 days and used to infect HEK293T cells followed by puromycin selection for 5 days. Knock-down efficiency was verified by Western blot and qPCR. qPCR primers were listed as follows: TRMT6-qFWD: CTGTCTTTGCTGGACTTTGTGGC; TRMT6-qRVS: AGACAGCCTGAGGTTGATGACC; TRMT61A-qFWD: TCCTCTACTCCACAGACATCGC; TRMT61A-qRVS: CAATGGTGCGGATGATGGCGTG.

### Quantification of m^1^A and m^6^A level by LC-MS/MS

200 ng isolated RNA or 100 ng model RNA oligo was digested into nucleosides by 0.5 U nuclease P1 (Sigma, N8630) in 20 μL buffer containing 10 mM ammonium acetate, pH 5.3 at 42 °C for 6 h, followed by the addition of 2.5 μL 0.5 M MES buffer, pH 6.5 and 0.5 U alkaline phosphatase (Sigma, P4252). The mixture was incubated at 37 °C for another 6 h and diluted to 50 μL. 5 μL of the solution was injected into LC-MS/MS. The nucleosides were separated by ultra-performance liquid chromatography on a C18 column, and then detected by triple-quadrupole mass spectrometer (AB SCIEX QTRAP 5500) in the positive ion multiple reaction-monitoring (MRM) mode. The mass transitions of m/z 282.0 to 150.1 (m^1^A), m/z 282.0 to 150.1 (m^6^A), m/z 268.0 to 136.0 (A) were monitored and recorded. Concentrations of nucleosides in RNA samples were deduced by fitting the signal intensities into the stand curves.

### Synthetic m^1^A RNA model sequences

Two pairs of synthetic m^1^A and A RNA oligoes were used in this study. The oligo sequences were listed as follows: m^1^A-1: CGCGGCUCG<u>m^1^A</u>GCCCGCGUGCGGGCCUCUUUCAGGCCGCU; A-1: CGCGGCUCG<u>A</u>GCCCGCGUGCGGGCCUCUUUCAGGCCGCU; m^1^A-2:CGGCGGCCCGGGACCG<u>m^1^A</u>GACCCGGCCCCGGCUCCCC; A-2:CGGCGGCCCGGGACCG<u>A</u>GACCCGGCCCCGGCUCCCC. The m^6^A contamination in the m^1^A oligoes caused during the oligo purification process was measured using quantitative LC-MS/MS. The m^1^A RNA oligoes and A RNA oligoes were mixed at the ratio: 100%, 75%, 50%, 25%,12.5%, 6.25% and 0%, respectively. The mixed m^1^A /A oligoes were subjected to library construction using specific RT primers as listed:

RT-m^1^A /A-1: ACACGACGCTCTTCCGATCTagcggcctgaaagaggc;
RT-m^1^A /A-2: ACACGACGCT CTT CCGAT CTggggagccggggcc.

### Cloning, Expression and Purification of AlkB

A truncated AlkB with deletion of the N-terminal 11 amino acids was cloned into pET30a (Novagen) and transformed to E. coli BL21(DE3) followed by growing in LB medium at 37 °C until the OD_600_ reached 0.6-0.8 and incubating at 30 °C for additional 4 h with the addition of 1 mM IPTG. Proteins were purified using Ni-NTA chromatography (GE Healthcare) and gel-filtration chromatography (Superdex 200, GE Healthcare) followed by Mono-Q anion exchange chromatography (GE Healthcare). Such purification procedure effectively avoided RNA contamination from *E coli.* (expression host).

### *In vitro* Demethylation treatment

*In vitro* demethylation treatment mediated by the demethylase AlkB: 10 μg full length polyA^+^ RNA was fragmented at 95 °C for 5 min using magnesium RNA fragmentation buffer (NEB) and fragmented polyA^+^ RNA was desalted and concentrated by ethanol precipitation. 10 μg (~0.2 nmol) fragmented polyA^+^ RNA was denatured at 65 °C for 5 min, and then demethylated in a 500 μL demethylation mixture containing 0.4 nmol purified AlkB, 50 mM MES, pH 6.5, 283 μM of (NH_4_)_2_Fe(SO_4_)_2_′6H_2_O, 300 μM 2-ketoglutarate, 2 mM L-ascorbic acid, 1 U/μL SUPERaseIn RNase Inhibitor (Invitrogen). The demethylation reaction was incubated for 2 h at 37 °C and quenched by the addition of 5 mM EDTA. The demethylated RNA was then purified by phenol chloroform extraction.

*In vitro* demethylation treatment mediated by the Dimroth rearrangement: 10 μg full length polyA^+^ RNA was incubated in alkaline buffer (0.1 M Na_2_CO_3_ /NaHCO_3_, 5mM EDTA, pH 10.2) at 65 °C for 3 h, and then the treated RNA was purified by ethanol precipitation.

### m^1^A-MAP

40 μg polyA^+^ RNA was fragmented into ~150 nt using magnesium RNA fragmentation buffer (NEB). m^1^A-containing RNA fragments were enriched by m^1^A immunoprecipitation as previous described (Li et al., 2016a). 10 ng (~0.2 pmol) of the immunoprecipitated m^1^A-containing RNA fragments were subjected to the AlkB demethylation treatment. RNA fragments were demethylated in a 20 μL demethylation mixture containing 0.4 pmol purified AlkB, 50 mM MES, pH 6.5, 283 μM of (NH_4_)_2_Fe(SO_4_)_2_·6H_2_O, 300 μM 2-ketoglutarate, 2 mM L-ascorbic acid, 0.4 U/μL RNase inhibitor and then incubated 2 h at 37 °C. The demethylation reaction was quenched by the addition of 5 mM EDTA, and demethylated RNA was purified by phenol chloroform extraction and ethanol precipitation.

Fragmented polyA^+^ RNA (as “input”), immunoprecipitated RNA [as (−) demethylase sample] and demethylated immunoprecipitated RNA [as (+) demethylase samples] were subjected to library construction. The library construction was performed according to the eCLIP library construction protocol with several modifications (Van Nostrand et al., 2016). For dephosphorylation of 3’ ends, RNA samples were treated with PNK (NEB) and incubated at 37 °C for 1 h and then heat-inactivation of PNK at 65 °C for 20 min. RNA samples were purified by ethanol precipitation and then subjected to 3’ RNA linker ligation using T4 RNA ligase2,truncated KQ (NEB) at 25 °C 2 h. The 3’ RNA linker sequence was listed: 5’rAPP-AGATCGGAAGAGCGTCGTG-3SpC3. The excess RNA adaptor was digested by adding 1 μL 5′ Deadenylase (NEB) into the ligation mix, incubating at 30 °C for 1 h and then adding 1 μL RecJf (NEB), incubating at 37 °C for another 1 h. These enzymes were subjected to heat-inactivation at 70 °C for 20 min and RNA was purified by ethanol precipitation. RNA pellets were dissolved in 10 μL H_2_O and then 1 μL 2 RT primer (ACACGACGCTCTTCCGATCT) was added. RNA-primer mix was denatured at 80 °C for 2 min and then chilling on ice. RT reaction buffer (50 mM Tri-HCl pH 8.3, 75 mM KCl, and 3 mM MgCl_2_, final), dNTPs (1 mM, final), DTT(5 mM, final), RNase Inhibitor (1 U/μL, final) and 1 μL TGIRT (InGex) were added into the denatured RNA-primer mix and reverse transcription was carried at 57 °C for 2 h. Excess RT primer was digested by the addition of 1 μL Exonuclease I (NEB) and incubation at 37 °C for 30 min. cDNA was purified using silane beads (Invitrogen) and then ligated to 5’ adaptor (5Phos-NNNNNNNNNNAGATCGGAAGAGCACACGTCTG-3SpC3). Ligation was performed with T4 RNA ligase 1, high concentration (NEB) at 25 °C overnight. The cDNA was purified using silane beads and then amplified by PCR with primers (5‘-AATGATACGGCGACCACCGAGATCTACACTCTTTCCCTACACGAC GCTCTTCCGATCT-3’; 5‘-CAAGCAGAAGACGGCATACGAGATXXXXXXGTGACTGGAGTTCAGACGTGTGCTCTTCCGA TC-3’, XXXXXX represents index sequence). PCR products were purified by 8% TBE gel and sequenced on Illumina Hiseq X10 with paired-end 2×150 bp read length.

### Locus-specific m^1^A detection

200 ng polyA^+^ RNA was isolated form TRMT6/61A stable knock-down cells and mock control cells respectively, and then were fragmented into 150 nt using magnesium RNA fragmentation buffer (NEB). Fragmented RNA was ligated to 3’ RNA linker and then reverse transcription was carried out with TGIRT using the same RT condition as that of m^1^A-MAP. The regions containing m^1^A were amplified by PCR using specific primes. And these amplicons from the same sample were mixed together and then subjected to DNA library construction using NEBNext^®^ Ultra™ II DNA Library Prep Kit for Illumina^®^ (E7645) according to the manufacturer’s instructions. These libraries were deep sequenced on Illumina Hiseq X10 with paired-end 2×150 bp read length. Thus, these regions of particular interest were covered with very high sequencing depth. Specific PCR primers were listed:

NM_025099-m^1^A5643-FWD: AAAAAGCTCGGTCCGGGTTC;
NM_025099-m^1^A5643-RVS: TTAGCCGCAAAATCACGCTG;
NM_001193375-m^1^A978-FWD: ACACCTGTCCAAGCCCTAAT;
NM_001193375-m^1^A978-RVS: CTGAGGGGCCCTTATTCCCA;
NR_026951-m^1^A659-FWD: ACGTCGGCTCGTTGGTCTAG;
NR_026951-m^1^A659-RVS: ACAGTCAAGCCTCCTGCAGC;
NM_001256443-m^1^A486-FWD: CAAGGTTCCAGGCGAAGGG;
NM_001256443-m^1^A486-RVS: AGCCGGGGTCTCTGTGG.

### siRNA Knock-down and overexpression of TRMT61B

Two synthesized duplex RNAi oligoes targeting TRMT61B mRNA sequences were used: 5‘-GGAUAUCAACCCAGGUGAUTT-3’ and 5‘-GCGUGAUUCAUGGAAAUUATT-3’ ; a scrambled duplex RNAi oligo (5’-UCCUCCGAACGUGUCACGUTT) was used as a mock control. The siRNA oligo was transfected into HEK293T by Lipofectamine RNAiMAX (Invitrogen) according to the manufacturer’s instructions and cells were harvested 48 h after transfection. Knockdown efficiency was verified by qPCR. qPCR primers were listed as follows: TRMT61B-qFWD: CAGGAGCAACCGAAGACAT; TRMT61 B-qRVS:ATATACAGCACATACACCACCAT; TRMT61B was cloned into pcDNA3.1 using the following primers: FWD: TCGCGAAACACTATGCTAATGGC; RVS: GTTAAGTTGTGGTTTGACCTTCCTC. The empty pcDNA3.1 vector was used as the mock control. The plasmid was transfected into HEK293T by Lipofectamine 2000 (Invitrogen) according to the manufacturer's instructions and cells were harvested 48 h after transfection.

### Ribosome profiling

2 Plates of 15-cm HEK293T cells were grown to 90% confluency; CHX was then added to the medium at a final concentration of 100 μg/mL for 7 min. The cells were then harvested and lysed with 1 mL lysis buffer (10 mM Tris-HCl, pH 7.4, 5 mM MgC1_2_, 100 mM KCl, 1 % Triton X-100, 2 mM DTT, 100 μg/mL CHX, 0.5 U/μL RNase inhibitor, 1×complete protease inhibitor). The cell lysates were centrifuged at 15,000g for 15 min and the supernatant was collected followed by measuring the 0D260 of cell lysates. 100 μL lysates were kept as input sample and 1 mL TRIzol was added to purify the RNA. 1 μL Micrococcal Nuclease (NEB) per 250D was added to the remaining cell lysates and allowed to incubate at 25 °C for 20 min. The digested cell lysates were used for performing ribosome foot-printing. Lysates were fractioned on 10/50% w/v sucrose gradients using the SW-40Ti rotor at 27,500rpm for 4h. 80S monosome fractions were collected followed by the addition of equal volume of extraction buffer (1% SDS, 40 mM EDTA). RNA was isolated by phenol-chloroform extraction. RNA fragments between 28-30 nt were selected using 15% Urea-PAGE. Recovered RNA fragments were subjected to library construction. In brief,RNA samples were dephosphorylated with PNK (NEB) and ligated to 3’ RP linker (5’rAPP-CTGTAGGCACCATCAAT-3SpC3) using T4 RNA ligase2, truncated KQ (NEB). Reverse transcription was carried using Superscript III (Invitrogen) with RP-RT primer (5Phos-AGAT CGGAAGAGCGT CGTGT AGGGAAAGAGTGTAGAT CTCGGTGGT CGC-SpC 18-CACT CA-SpC18-TTCAGACGTGTGCTCTTCCGATCTATTGATGGTGCCTACAG). cDNA was circ-ligated with CircLigase II (Epicentre) and then amplified by PCR with primers (5‘-AATGATACGGCGACCACCGAGATCTACAC-3’; 5‘-CAAGCAGAAGACGGCATACGAGATXXXXXXGTGACTGGAGTTCAGACGT GTGCT CTTCCGA TC-3’, XXXXXX represents index sequence). PCR products were purified by 8% TBE gel and sequenced on Illumina Hiseq 2500 with single end reads (50 bp).

### qPCR-based m^1^A level evaluation

20 μg polyA^+^ RNA was isolated from HEK293T cells with TRMT61B overexpression, knock-down and the corresponding mock controls, respectively. RNA was fragmented into ~150 nt using magnesium RNA fragmentation buffer (NEB) and concentrated by ethanol precipitation. Fragmented RNA (as input) was denatured and incubated with 2 μg anti-m^1^A antibody in IPP buffer (150 mM NaCl, 0.1% NP-40, 10 mM Tris, pH 7.4) at 4°C overnight. 20 μl Protein A/G UltraLink Resin (Pierce) was added to the RNA antibody mixture and incubated for additional 3 h at 4°C. Resins were washed with twice with IPP buffer, once with low salt buffer (75 mM NaCl, 0.1% NP-40, 10 mM Tris, pH 7.4), once with high salt buffer (200 mM NaCl, 0.1% NP-40, 10 mM Tris, pH 7.4) and twice with TEN buffer (10 mM Tris, pH 7.4, 1 mM EDTA, pH 8.0, 0.05% NP-40). m^1^A-containing RNA was eluted from resins with 3 mg/mL *N*^1^-methyladenosine (Berry&Associates) in IPP buffer and purified by phenol chloroform extraction and ethanol precipitation. Input and immunoprecipitated RNAs were reverse transcribed into cDNA using Superscript III (Invitrogen) and quantified by qPCR using SYBR GREEN mix (Takara) on Roche Lightcycler 96 real-time PCR system. The m^1^A-IP/input ratio of target regions in the TRMT61B overexpression, knock-down and the corresponding control samples were calculated, respectively. The primers used for qPCR were listed below:

MT-CO1-qFWD: CCTATCATCTGTAGGCTCATTC;
MT-CO1-qRVS: GGAGGGTTCTTCTACTATTAGGAC;
MT-CO2-qFWD: ACAGATGCAATTCCCGGACG;
MT-CO2-qRVS: CCACAGATTTCAGAGCATTGACC;
MT-CO3-qFWD: CGCCTGATACTGGCATTTTG;
MT-CO3-qRVS: GACCCTCATCAATAGATGGAGAC;
MT-CYB-qFWD: CAACCCCCTAGGAATCACCTC;
MT-CYB-qRVS: GAGGGCGTCTTTGATTGTGTAG;
16S rRNA-qFWD: ATGAATGGCTCCACGAGGG;
16S rRNA-qRVS: CTTGCTGTGTTATGCCCGC.

### Reductive dimethylation labeling

Mitochondria was isolated from TRMT61B overexpression and mock control HEK293T cell lines according to the manufacturer’s (Thermo Fisher) and lysed with RIPA buffer followed by sonication. After extraction, total proteins from different cell lines were quantified with the BCA protein assay kit (Thermo Fisher). Equal amount of proteins from two cell lines were digested by trypsin on-column in 100 mM TEAB buffer and subjected to reductive dimethylation labeling. 4 μL of 4% (w/w) light or heavy formaldehyde was added to 100 μL of trypsin digested samples prepared from TRMT61B overexpression and mock control HEK293T cells, respectively. In the meantime, 4 μL of 0.6 M sodium cyanoborohydride was added and the samples were incubated at room temperature for 1h. The dimethylation labeling reaction was quenched by the addition of 1% (w/w) ammonia and 5% (w/w) formic acid. Finally, light and heavy labeled peptide samples were mixed, concentrated by vacuum, and analyzed on a Q Exactive mass spectrometer (Thermo Fisher).

### LC-MS/MS and data analysis

The peptides were analyzed on a Q Exactive mass spectrometer (Thermo Fisher). Under the positive-ion mode, full-scan mass spectra was acquired over the m/z range from 350 to 1800 using the Orbitrap mass analyzer with mass resolution setting of 70000. MS/MS fragmentation was performed in a data-dependent mode, of which the 20 most intense ions were selected from each full-scan mass spectrum for high-energy collision induced dissociation (HCD) and MS2 analysis. MS2 spectra were acquired with a resolution setting of 17500 using the Orbitrap analyzer. Some other parameters in the centroid format: isolation window, 2.0 m/z units; default charge, 2+; normalized collision energy, 28%; maximum IT, 50 ms; dynamic exclusion, 20.0 s.

LC-MS/MS data was analyzed by ProLuCID (Xu et al., 2015) with static modification of cysteine (+57.0215 Da) and variable oxidation of methionine (+15.9949 Da). The searching results were filtered by DTASelect (Tabb et al., 2002) and peptides were also restricted to fully tryptic with a defined peptide false positive rate of 1%. The ratios of reductive dimethylation were quantified by the CIMAGE software as described before (Weerapana et al., 2010).

### Pre-processing of raw sequencing data

A random barcode of 10 nt was included in the adapter that ligates to the 3’ end of cDNA and it cannot be precisely located in Read 1. Hence, only Read 2 data of m^1^A-MAP were used for subsequent analyses. Raw sequencing reads produced from m^1^A-MAP were firstly subjected to Trim_galore (http://www.bioinformatics.babraham.ac.uk/projects/trim_galore/) for quality control and adaptor trimming. The minimum quality threshold was set to 20, and the minimum length required for reads after trimming was 30 nt. The remaining reads were further processed by removing the first 10 nt random barcode in the 5’ end. As for the ribosome profiling (Ribo-seq) data and the corresponding RNA-seq data, reads with a quality lower than 20 were discarded, and the adaptor in 3’ end was trimmed. Processed reads with a length ranging from 25 nt to 35 nt in the ribosome profiling sample were kept for further analysis.

### Reads mapping and PCR duplication removing

Processed reads were mapped to human transcriptome or mitochondrial genome using BWA-MEM with default parameters (version 0.7.15-r1140) (Li and Durbin, 2009). Reference transcriptome was prepared based on the Refseq annotation of human (hg19) downloaded from the table browser of UCSC database. The redundant sequences with the same Refseq id were removed. Transfer RNA (tRNA) sequences were also downloaded from UCSC table browser and integrated into the transcriptome. Mitochondrial genome and corresponding annotation were downloaded from NCBI (NC_012920.1). Reads mapping to an identical position of reference were considered as PCR duplications if their 10 nt random barcodes were the same, and only one of these reads was kept. Performances related to the processing of sam/bam file were done with the help of SAMtools (Li et al., 2009) (http://samtools.sourceforge.net/).

### Identification of m^1^A sites

Mismatch rate of each nucleotide in the reference sequences was calculated for both (−) and (+) demethylase samples. Two parameters were defined to evaluate the dynamic change of mismatch rate in the (−) demethylase and (+) demethylase samples: difference of mismatch rate (Diff) and fold change of mismatch rate (FC). Diff was calculated by subtracting the mismatch rate in the (+) demethylase from that in the (−) demethylase samples, while FC was calculated by dividing the mismatch rate in the (−) demethylase by that in the (+) demethylase sample. FC was artificially set to “1000” if the mismatch rate in the (+) demethylase sample was “0”. A position was identified as an m^1^A site when the following criteria were met: a) FC>=3; b) Diff>=10%; c) the number of reads with a mismatch at the position was no less than 5; d) criteria (a-c) were all fulfilled in both replicates.

In order to evaluate the frequency of potential false positives caused by m^1^A-independent mismatch, we employed a reverse calling procedure. Specifically, the “opposite” calculations of Diff and FC values for each position were performed:

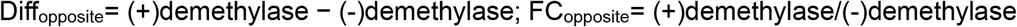

Under such circumstance, only 17 sites passed the above-mentioned threshold, suggesting the m^1^A-independent mismatch should minimally interface with the identification of true m1A sites.

### Motif discovery and GO enrichment analysis

For the analysis of sequence consensus, 15 nt of sequence neighboring each m^1^A site was retrieved. These sequences were then subjected to DREME algorithm in MEME suite (Version 4.12.0) for the discovery of enriched motifs (Bailey et al., 2009). The shuffled input sequences were used as the background to eliminate potential false positives caused by the nucleotide composition.

Gene Ontology (GO) enrichment analyses were performed using DAVID web-based tool (http://david.abcc.ncifcrf.gov/) (Huang da et al., 2009).

### Secondary structure and minimum free energy analysis

12 nucleotides of the 5’ end and 10 nucleotides of the 3’ end of each m^1^A site (hence 23 nt sequence in total) were retrieved for local structure analysis. RNAfold (v2.3.4) (http://rna.tbi.univie.ac.at/cgi-bin/RNAfold.cgi) was used to predict the secondary structure and calculate the corresponding minimum free energy (MFE). The length of the loop where m^1^A resides was determined based on the predicted structure; for an m^1^A site that is not located in a loop, this value was set to “0”. The significance test of MFE between m^1^A sites within the GUUCRA motif and other m^1^A sites was performed using Mann-Whitney U-test.

### Ribosome profiling data analysis and TE calculation

Ribo-seq and corresponding RNA-seq reads were aligned to the transcriptome, and RPKM (Reads Per Kilobase per Million mapped reads) of each transcript was calculated. Translation efficiency (TE) was defined for each transcript as the ratio of RPKM in Ribo-seq to RPKM in RNA-seq.

For the analysis of influence of m^1^A on mitochondrial gene translation, mitochondrial ribosome profiling data were downloaded from the GEO Datasets (GSE48933) (Rooijers et al., 2013). The depth of reads covered for each nucleotide along the mitochondrial transcripts was retrieved using Samtools depth tool.

### miRNA target analysis

The predicted miRNA targeting sites on mitochondrial coding genes were downloaded from the miRWalk database (v2.0) (Dweep and Gretz, 2015), which depends on the match of “seed region” to gene sequence. The minimum length required for the match of seed region was set to 7 nt. The experimentally identified miRNA targeting sites were retrieved from the published CLASH results (Helwak et al., 2013).

